# Sustained Yap/Taz activation promotes aberrant alveolar epithelial cell differentiation and drives persistent fibrotic remodeling

**DOI:** 10.1101/2025.07.16.665213

**Authors:** Isabella P. Gaona, A. Scott McCall, Natalie M. Geis, Arlo C. Colvard, Gianluca T. DiGiovanni, Taylor P. Sherrill, Ujjal K. Singha, David S. Nichols, Anna P. Serezani, Holly E. David, Jean-Philippe Cartailler, Shristi Shrestha, Sergey S. Gutor, Timothy S. Blackwell, Jonathan A. Kropski, Jason J. Gokey

**Affiliations:** Division of Allergy, Pulmonary, and Critical Care Medicine, Department of Medicine, Vanderbilt University Medical Center, Nashville, TN 37232, USA; Department of Veterans Affairs Medical Center, Nashville, TN 37212, USA; Vanderbilt University, Nashville, TN 37232, USA; Department of Internal Medicine, University of Michigan Medical School, Ann Arbor, MI 48109, USA; Department of Cell and Developmental Biology, Vanderbilt University School of Medicine, Nashville, TN 37232, USA

## Abstract

YAP/TAZ signaling is required for initiation of lung alveolar repair, yet previous studies in idiopathic pulmonary fibrosis (IPF) predicted increased YAP/TAZ signaling in alveolar epithelial cells (AECs). We investigated whether persistent YAP/TAZ AEC signaling contributes to failed epithelial repair and persistent fibrotic remodeling. In IPF lungs, we identified increased YAP^+^/TAZ^+^ AECs and increased expression of YAP/TAZ transcriptional targets compared to donor control lungs. In human lung organoids, pharmacological YAP/TAZ activation resulted in phenotype shifts of AECs into aberrant transitional states. In mice with Yap/Taz activation (YT^active^) resulting from deletion of Hippo-kinases Stk3/4 in alveolar-type 2 (AT2) cells, resulted in persistent fibrotic remodeling at 28- and 56-days post-bleomycin injury. Gene promoter activity associated with transitional cell markers (*Krt19, Hopx,* and *Runx2*) was increased in YT^active^ AT2 cells. Immunofluorescent staining showed a loss of AT2 associated Cebpa and increased *Krt19* in YT^active^ lineage traced AT2 cells 28 days post-injury. Inhibition of Yap/Taz using Verteporfin resulted in improved lung repair in YT^active^ mouse lungs, including increased Cebpa and decreased *Krt19^+^* transitional cells. These findings demonstrate sustained Yap/Taz activation drives abnormal alveolar repair and persistent fibrotic remodeling. Blocking aberrant persistent Yap/Taz activity promotes adaptive repair and has potential as a therapeutic strategy for PF.

## Introduction

Idiopathic Pulmonary Fibrosis (IPF) is a chronic and progressive lung disease characterized by the replacement of functional alveoli with dense fibrotic scarring^1,2^. This fibrotic remodeling is associated with loss of respiratory function and is often lethal within 3-5 years of diagnosis. The only 2 FDA approved treatment for IPF modestly slow loss of respiratory function, but do not stabilize disease or improve quality of life for IPF patients^3–5^. The cause of IPF is unknown, but evidence from genetic studies and mouse models indicate that repetitive injury to – and failed repair of - the alveolar epithelium is a driving factor of disease progression that leads to activation of the fibroblast^6–10^.

In the human and mouse lung alveolar epithelium, the alveolar type 2 (AT2) cells act as facultative progenitor cells during alveolar repair, proliferating to generate more AT2 cells and differentiating into alveolar type 1 (AT1) cells that lay in close proximity to the alveolar capillaries to facilitate gas exchange^11–14^. During “failed” alveolar repair, aberrant progenitor AT2 populations can form “hyperplastic AT2” regions or differentiate into persisting alveolar transitional cell populations expressing markers of both AT1/AT2 cells that do not adopt the long thin AT1 cell shape that is essential to facilitate gas exchange^15–18^. These abnormal repair processes that are believed to be an early driver of pulmonary fibrosis, thus understanding the mechanisms that drive normal “adaptive” repair versus those that lead to fibrotic remodeling is essential to develop treatments that will halt disease progression and/or restore lung function.

Several developmental pathways that are typically quiescent during lung homeostasis become transiently activated during normal repair, however persistent activation of these pathways is predicted to lead to aberrant injury responses^15,16,19,20^. Recent work has implicated the Hippo-Yap/Taz pathway being involved in AT2 to AT1 differentiation during development and repair^21–23^. The Hippo-Yap/Taz pathway consists of the Hippo components mammalian serine-threonine kinases 1/2 (MST1/2 in humans, Stk4/3 respectively in mouse) that phosphorylate Lats1/2 that in turn phosphorylate Yap and Taz. When this phosphorylation cascade is active, Yap and Taz are sequestered in the cytoplasm, however in the absence of phosphorylation Yap and/or Taz translocate to the nucleus where they interact with DNA binding partners^24–26^ (classically TEADs^27–29^ and also with Runx2^30,31^ and Smads^32–34^) to direct transcription of target genes associated with proliferation and differentiation. Previous work from our group and others determined that activation of Yap/Taz signaling is required for alveolar repair, as deletion of Yap/Taz or just Taz in AT2 cells prevents AT1 cell differentiation^35–38^. Our previous work demonstrated nuclear Taz is expressed in nearly all AT1 cells, while nuclear Yap or Taz is rarely detected in AT2 cells at homeostasis. During injury repair, Yap and Taz are activated in AT2 cells, with peak activity at 7 and 14 days respectively, and are down-regulated to low activity by 21-28 days post-bleomycin injury^35^. These studies demonstrate that some degree of Yap/Taz activity is essential early during the repair process and preemptive inhibition of Yap/Taz would be maladaptive for the lung epithelium. In contrast, scRNA-sequencing and immunofluorescence data from our group and others have predicted that increased YAP/TAZ activity is associated with aberrant epithelial cells in the IPF lung as well as fibroblast activation^37,39–45^. This led us to hypothesize that aberrant and persistent Yap/Taz activation in AT2 cells prevents normal alveolar epithelial repair, results in persistent transitional cells, and promotes fibrotic remodeling in the lung. In these studies, we sought to define the role of sustained Yap/Yaz activity in AT2 cells during lung injury and subsequent fibrotic remodeling and test whether interrupting this persistent epithelial Yap/Taz activity would promote adaptive repair and block or delay IPF progression.

## Results

To determine whether YAP and/or TAZ is activated in the IPF lung epithelium, we first performed immunofluorescence staining to assess nuclear YAP or TAZ in SP-C^+^ (either marking AT2 cells in healthy donor/IPF or hyper/metaplastic alveolar epithelium in IPF) cells or AGER^+^ (labeling AT1 in healthy donor/IPF or aberrant epithelium in IPF) cells. De-identified donor and IPF subject (N=6 each) demographics are reported in supplemental **Table S1**. Consistent with our previous findings^41^, YAP was rarely detected in SP-C^+^ cells (5.8^+^/-4.2%) of healthy donors, however YAP was present in the nucleus of 38.7% (^+^/-6.3%) of SP-C^+^ cells including cells that were both SP-C^+^/AGER^+^ in IPF lungs (**Figure 1A, C**). TAZ was also rarely detected (2.1^+^/-2.4%) in healthy donor SP-C^+^ AT2 cells, however nuclear TAZ was present in 21.4% (^+^/-7.3%) of SP-C^+^ and SP-C^+^/AGER^+^ cells in IPF lungs (**Figure 1B, D**). We then interrogated scRNA-seq data from our recently published work^46,47^, performing cell-type specific, subject-level pseudobulk differential expression analysis comparing IPF and control AT2 cells. We found that *YAP* and *TAZ* (*WWTR1*), YAP/TAZ binding partners (*TEAD1/3, NFIB*), and YAP/TAZ transcription targets were increased in IPF AT2 cells compared to control donor AT2 cells (**Figure 1E)**. These findings support the concept that there is sustained activation of Hippo-YAP/TAZ signaling in IPF alveolar epithelium.

**Figure 1.**
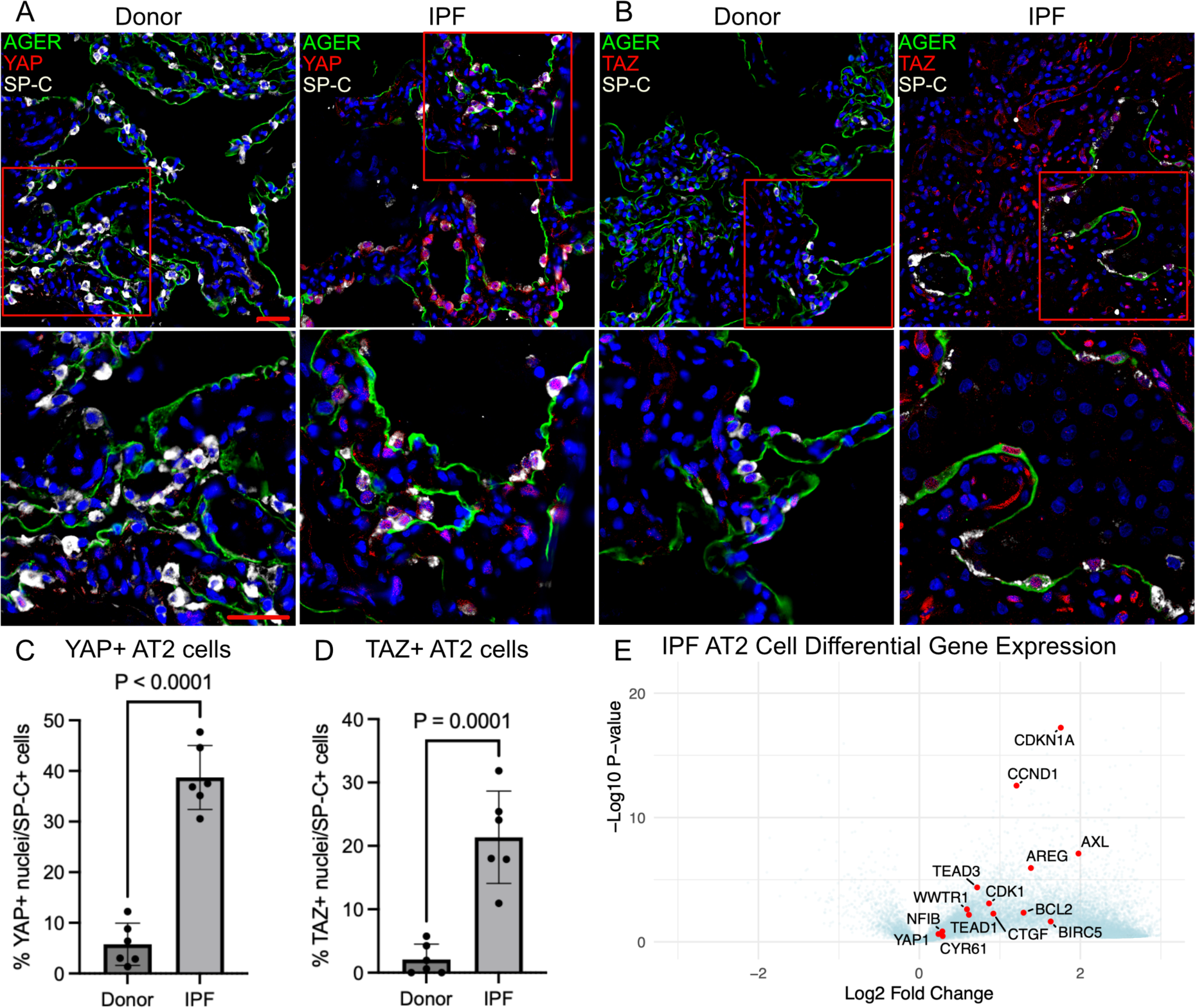
YAP and TAZ are active in the IPF epithelium. **A**) Immunofluorescence analysis of YAP (red) and (**B**) TAZ (red) in AGER^+^ (green) AT1 cells and SP-C^+^ (white) AT2 cells. **C**) Quantification of YAP^+^ nuclei and (**D**) TAZ^+^ nuclei in SP-C^+^ AT2 cells from donor and IPF lungs. *n* = 6 per group. Statistics determined using an unpaired parametric t-tests. **E**) Differential gene expression analysis of IPF AT2 cells generated from previously published scRNAseq analysis of IPF and donor lungs (GSE227136). Scale bar represents 50μm.

To test the consequences of sustained YAP/TAZ signaling in human alveolar epithelial cells, we isolated CD326^+^ epithelial cells from peripheral lung tissue obtained from declined lung donors and cultured them serum-free-feeder-free media as alveolar organoids^48^. Organoids were grown from day 1-7 in SFFFM media and then either continued in SFFFM until day 14 or switched to alveolar differentiation media from day 7-14 and were treated with either vehicle or the Yap/Taz activating drug XMU-MP1 (3μM), a specific MST1/2 inhibitor^49^, from culture day 4-14, in which we observed led to dramatic changes in organoid appearance (**Figure 2A)**. We noted morphologic differences in culture and thus sought to characterize multiple subcellular components to quantify cellular morphologies. Whole-mount staining of RNA (SYTO 14) cytoskeletal components (Wheat Germ Agglutinin and Phalloidin) dyes followed by high-content imaging and analysis were used^50^. We found that activation of YAP/TAZ with XMU-MP1 resulted in marked and over-riding morphologic changes of these human AT2 cell organoids whether cultured in SFFFM or commercial ADM (alveolar differentiation media) which drove unique morphologic appearance (ADM resulting in a splayed appearance) (**Figure 2B**). Analysis of vehicle and XMU treated organoids revealed cell morphology changes in the YAP/TAZ activated organoids as demonstrated by visualizing the variance in cell shape in phenotypic PCA space derived from all extracted morphologic features with significant shifts between SFFFM, ADM, or XMU treated organoids^51^ (**Figure 2C).** Likewise, analysis of cell nuclei morphology in phenotypic space incorporating RNA nuclear localization revealed significantly altered nuclear features in YAP/TAZ activated organoids which is consistent with prior YAP/TAZ activation phenotypes observed in mechanical situations^44,52–54^ (**Figure 2D).** We then further characterized these organoids using immunofluorescent staining and scRNA sequencing. Collected organoids were stained for AGER and SP-C which demonstrated that YAP/TAZ activation increased the expression of AGER especially in SFFFM organoids (**Figure 3A**). Analysis of scRNA data indicated that sustained activation of YAP/TAZ signaling resulted in the aberrant differentiation of AT2 cells into several sub-populations including a YAP/TAZ-active aberrant population expressing high levels of the mitochondrial marker *TOMM20* and a YAP/TAZ-active population expressing high levels of keratins (KRT^hi^) associated with aberrant transitional cell populations (**Figure 3B, C**). These aberrant populations were present in XMU-MP1 treated cultures regardless of the presence of SFFFM or ADM culture media (**Figure 3D, E)**, indicating that sustained activation of YAP/TAZ in human epithelial cells results in abnormal alveolar cell behavior and differentiation into aberrant cell populations which supersedes signaling provided by media with either AT2 or components of alveolar differentiation.

**Figure 2.**
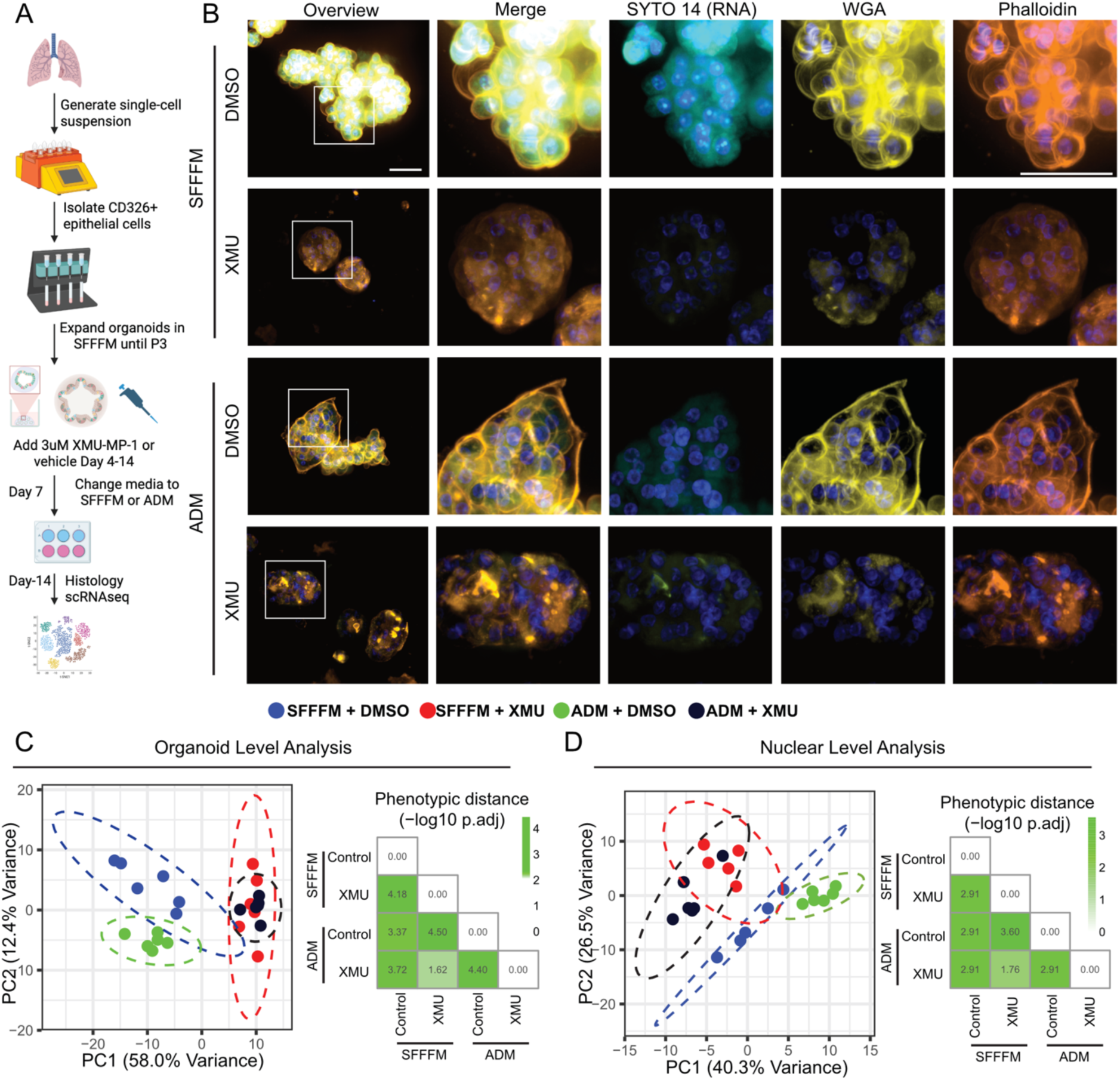
YAP/TAZ activation alters cell morphology in human organoids. **A**) Schematic of human epithelial organoid generation, YAP/TAZ activation (XMU-MP-1), and collection strategy. **B**) Cell and nuclei shape analysis using WGA (yellow) and Phalloidin (orange) with SYTO14 (cyan) in organoids treated with Vehicle or XMU-MP1 in either SFFFM or ADM. **C)** Organoid morphologic analysis across phenotypic PCA space based on analysis of high-dimensional feature extraction and statistical analysis of phenotypic between groups. Each point represents wells as technical replicates **D)** Analysis of cell nuclei across phenotypic PCA space and phenotypic distance between groups. Each point represents wells as technical replicates. P values adjusted for multiple comparisons. Scale bars represent 50μm.

**Figure 3.**
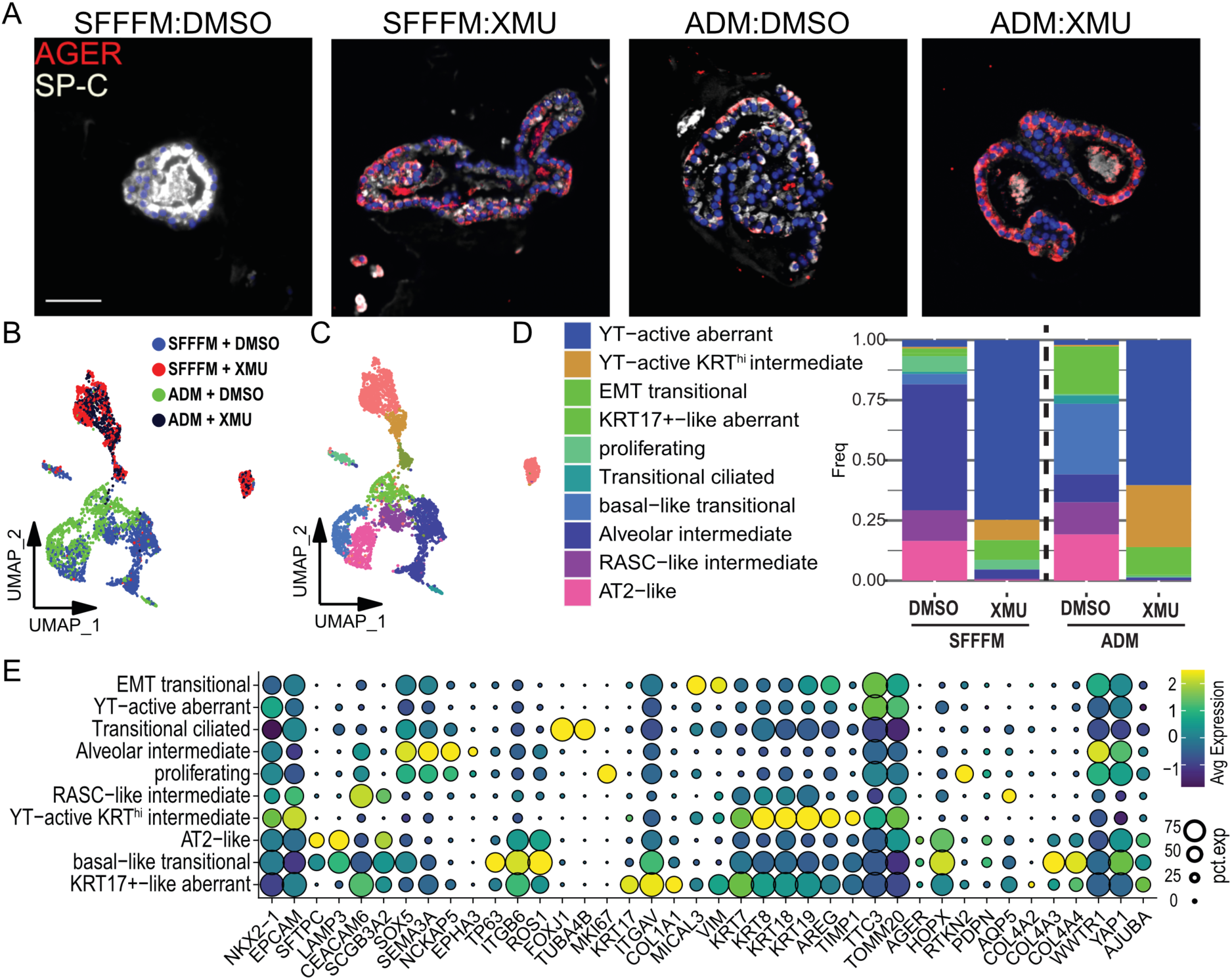
YAP/TAZ activation leads to aberrant epithelial cell populations in human alveolar organoids. **A**) Immunofluorescence analysis of AGER^+^ cells and SP-C^+^ cells in organoids treated with 3µM XMU-MP-1 or DMSO in expansion (SFFFM) and differentiation (ADM) media. **B-C**) UMAP embedding of jointly analyses scRNA-seq of YAP/TAZ activated and DMSO control organoids annotated by treatment condition (B) or cell type/state (C). **D**) Quantification of percentages of normal and aberrant cell populations. **E**) Dot plot depicting marker gene expression in normal and aberrant cell populations. Scale bars are 50μm.

Next, to test the impact of AT2-cell Yap/Taz activation on experimental lung fibrosis in-vivo, we turned to a mouse model of conditional Yap/Taz activation in AT2 cells. In these mice, Hippo-kinase floxed mice (*Stk3^fl/fl^/4^fl/fl^*) were crossed with *S*ftpcCre*^ert^*^2t^*^dTomato^*mice, and Cre^+^ Yap/Taz active mice (hereafter referred to as YT^active^ mice) and Cre^-^ littermate controls or *S*ftpcCre*^ert^*^2t^*^dTomato^* WT mice were administered tamoxifen (100mg/kg intraperitoneal) 3-weeks prior to a single intratracheal dose of saline or 0.08IU bleomycin. Lungs were collected at 14-days and 28-days post bleomycin for assessment of fibrosis (**Figure 4A-B**). While total collagen (**Figure 4C**) and Ashcroft scoring (**Figure 4D**) were similar in WT and YT^active^ on day 14 after bleomycin, by day 28 YT^active^ mice had increased Ashcroft score and lung collagen content compared to WT mice. This indicated that sustained Yap/Taz activation in AT2 cells drives persistent lung fibrosis and suggested the effects were mediated through regulation of the repair/resolution phase. To specifically test whether YT activation in AT2 cells during repair/resolution exacerbates fibrosis, we administered bleomycin to YT^active^ mice and littermate controls, then on day 14 post-bleomycin, mice were administered tamoxifen to delete *Stk3/4 (*thus activate YT) or corn oil (control) (**Figure 4E**). YT^active^ mice had increased collagen content (**Figure 4F**) and worsened Ashcroft score (**Figure 4G**) at both day 28 and day 56 post-bleomycin. Together, these results indicated that persistent activation of YT in AT2 cells prevented resolution of lung injury and resulted in persistent lung fibrosis.

**Figure 4.**
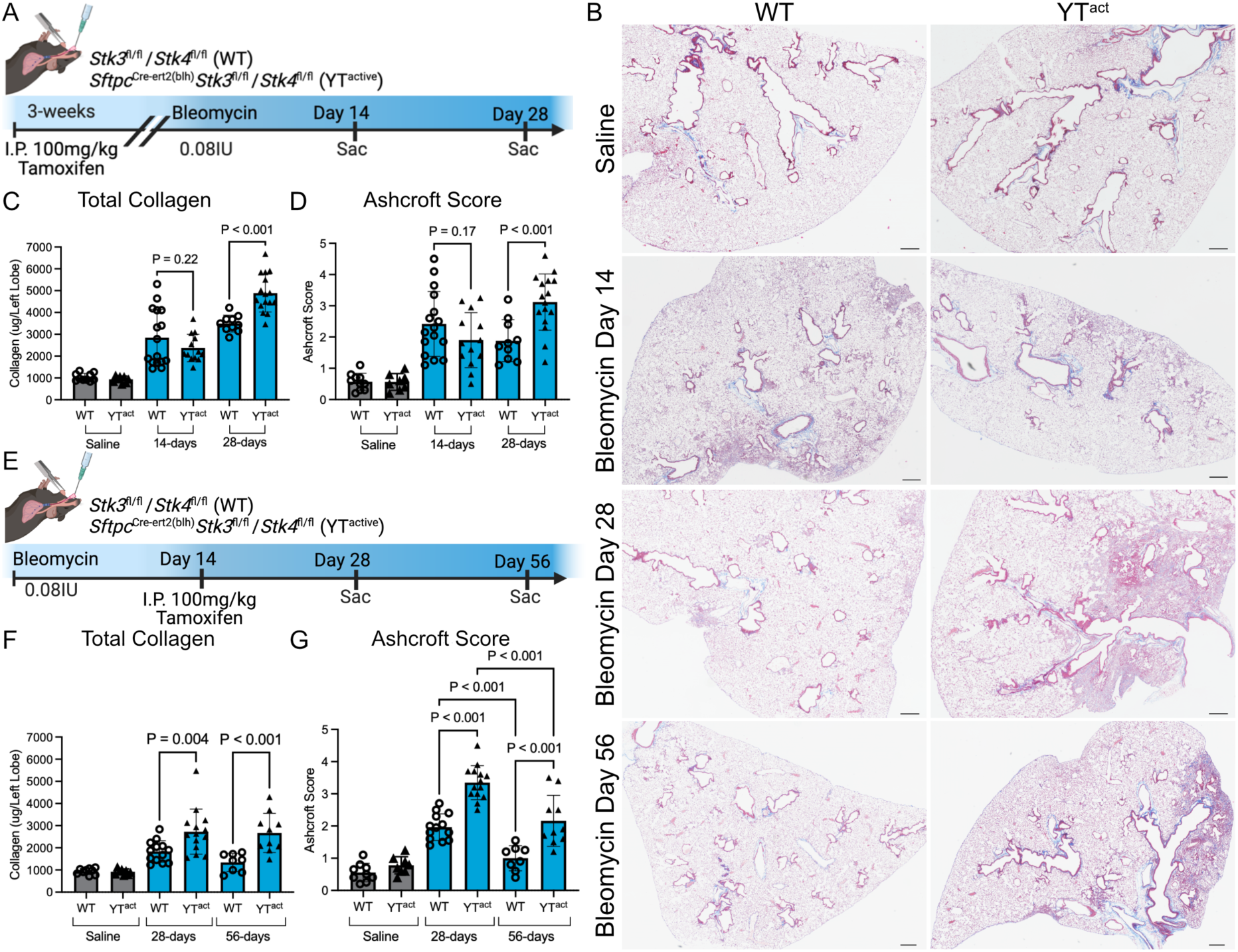
YT^active^ mice show increased and sustained fibrotic remodeling. **A**) Timeline of mouse injury model with Yap/Taz activated prior to bleomycin with lungs assessed at 14- and 28-days. **B**) Masson’s trichrome staining of lung tissue sections from wild-type and YT^active^ mice at 14-, 28- and 56-days after bleomycin or saline instillation. **C**) Quantitative analysis of total collagen and (**D**) Ashcroft scoring of fibrosis in WT and YT^active^ mice at 14- and 28-days after saline or bleomycin. WT saline (n=10), YT^active^ saline (n=8), day-14 WT bleomycin (n=16) and YT^active^ bleomycin (n=13), day-28 WT bleomycin (n=10) and YT^active^ (n=16) bleomycin treated mice. **E**) Timeline of mouse lung injury model with Yap/Taz activated after bleomycin and collection at 28- and 56-days. **F**) Quantitative analysis of total collagen and (**G**) Ashcroft scoring of fibrosis in wild-type and YT^active^ mice at 28- and 56-days after saline or bleomycin. WT saline (n=9), *n* = 8 YT^active^ saline (n=8), day-28 WT (n=13) and YT^active^ (n=14) bleomycin, day-56 WT (n=8) and YT^active^ (n=10) bleomycin-treated mice. Ordinary one-way ANOVA with Sidak’s multiple comparisons test with a single pooled variance was used to compare groups. Scale bars are 100μm.

Next, to determine if intervention with a pharmacological Yap/Taz inhibitor could restore alveolar repair/regeneration, tamoxifen-treated YT^active^ and WT mice were challenged with IT bleomycin and administered Verteporfin (60mg/kg intraperitoneal), a Yap/Taz-Tead inhibitor that prevents Yap/Taz nuclear localization and enhances degradation^55^, 14 days after bleomycin induced lung injury (**Figure 5A**). Consistent with the above findings, YT^active^ mice had increased collagen content and worsened lung injury scores at 28-days post-injury (**Figure 5B-D**). While Verteporfin did not impact fibrosis in WT mice, Verteporfin treatment of YT^active^ mice led to improved Ashcroft scores and lower total collagen content as measured by total sircol compared to vehicle-treated YT^active^ mice at 28-days post-injury (**Figure 5B-D**). Immunofluorescence analysis of WT and YT^active^ mouse lungs treated with saline or bleomycin showed increased Hopx^+^ AT1-like cells derived from lineage-labeled AT2 cells, as well as increased Sp-C^+^/Hopx^+^ transitional cells in YT^active^ bleomycin treated lungs (**Figure 5E-G**). Verteporfin treatment did not significantly impact the number of lineage-traced Hopx^+^ cells, however the number of transitional Sp-C^+^/Hopx^+^ cells was reduced, suggesting resolution of the transitional phenotype. YT^active^ bleomycin treated lungs had decreased overall Sp-C^+^ AT2 cells and increased Hopx^+^ AT1 like cells, which were not significantly affected by Verteporfin treatment (**Figure 5I, Supplemental Figure 1**). We observed that during single-dose bleomycin treatment, a small number of Scgb1a1^+^ airway cells (2.0^+^/-1.1%) migrate into the distal lung to potentially participate in repair. Scgb1a1^+^ cells were increased (10.3^+^/-6.2%) in the distal lung parenchyma in YT^active^ bleomycin-injured lungs compared to littermates, while Verteporfin treatment led to reduced parenchymal Scgb1a1^+^ cells (5.0^+^/-4.2%) compared to vehicle-treated YT^active^ mice (**Figure 5H, J**). These findings indicate that Verteporfin treatment after onset of fibrotic remodeling improved lung regeneration by antagonizing the sustained Yap/Taz activity in AT2 cells.

**Figure 5.**
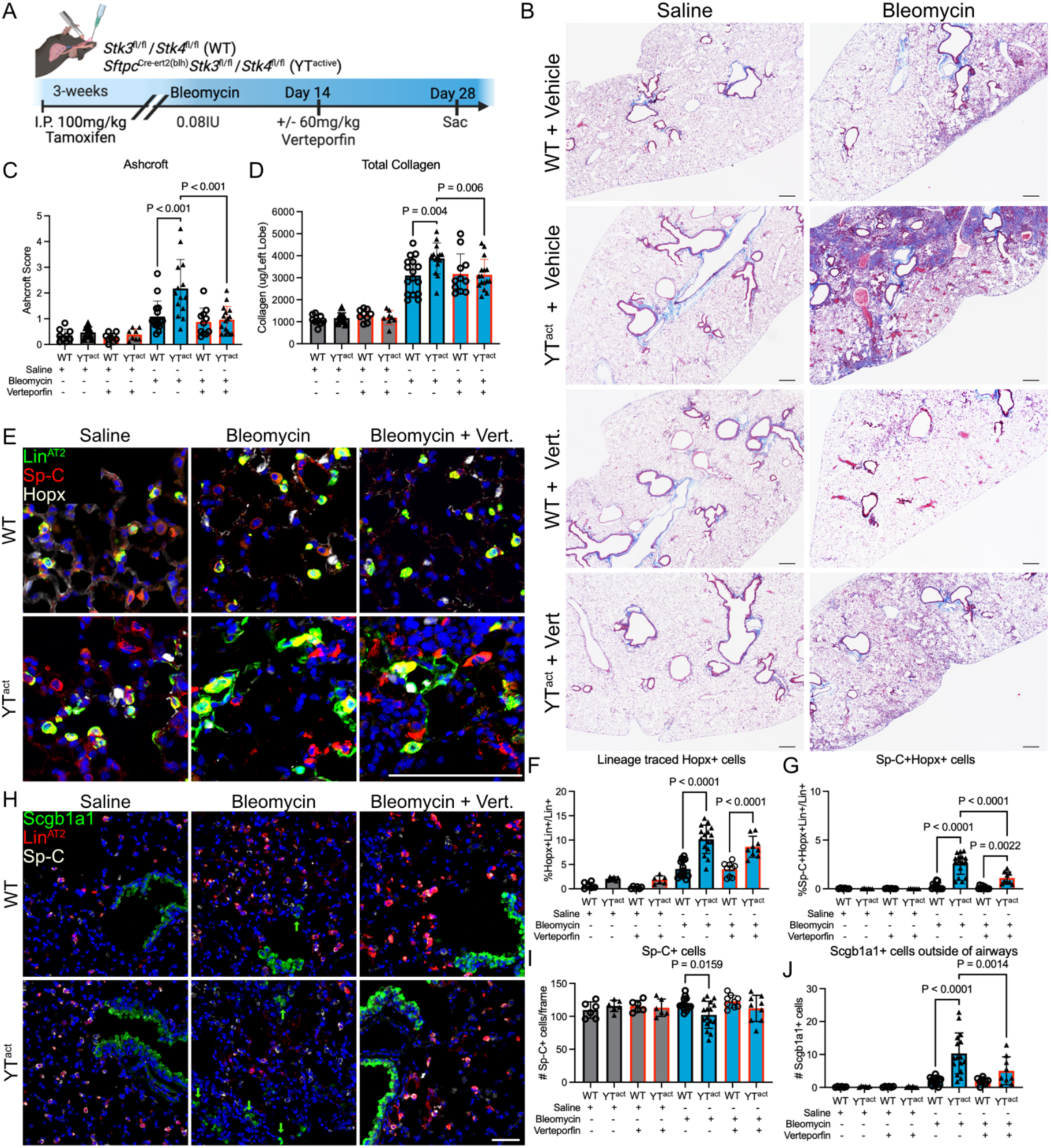
The Yap/Taz inhibitor Verteporfin partially rescues fibrotic phenotype in YT^active^ mice after bleomycin injury. **A**) Timeline of YT^active^ mouse bleomycin injury model with Yap/Taz inhibition with Verteporfin (60mg/kg) at 14-days post-injury. **B**) Masson’s trichrome staining of WT and YT^active^ mice with and without Verteporfin following saline or bleomycin. **C**) Ashcroft scoring of fibrosis and (**D**) total collagen analysis of wild-type and YT^active^ mice with and without Verteporfin following saline or bleomycin. WT saline (n=9). and YT^active^ saline (n=9), WT (n=9) and YT^active^ (n=8) saline and Verteporfin, WT (n=16) and YT^active^ (n=14) bleomycin, WT (n=11) and YT^active^ (n=16) bleomycin and Verteporfin-treated mice. **E**) Immunofluorescence analysis of Sp-C^+^ (red), Hopx^+^ (white), and lineage-traced AT2 (green) cells. Quantification of (**F**) lineage traced AT2 cells expressing Hopx, (**G**) Sp-C^+^/Hopx^+^ cells, and (**I**) total Sp-C^+^ cells per frame. **H**) Scgb1a1^+^ cells (green), AT2 lineage-labeled cells (red), and Sp-C^+^ cells (white) in wild-type and YT^active^ mice given saline, bleomycin, or bleomycin and Verteporfin. **J**) Quantification of Scgb1a1^+^ cells outside of airways in the alveolar region. WT and YT^active^ saline (n=6), WT and YT^active^ saline/Verteporfin (n=6), WT (n=14) and YT^active^ (n=16) bleomycin, WT (n=8) and YT^active^ (n=9) bleomycin/Verteporfin-treated mice. To determine significance an ordinary one-way ANOVA with Sidak’s multiple comparisons test with a single pooled variance was used. Immunofluorescent analysis scale bars represent 50μm. Trichrome staining scale bars are 100μm.

To examine the mechanisms through which sustained Yap/Taz activation in AT2 cells prevents functional alveolar repair and fibrosis resolution, we performed single-nucleus multiome (RNA^+^ATAC sequencing) of lung tissue from tamoxifen-treated WT and YT^active^ mice 28 days after bleomycin (or saline control). Following data integration, cells were clustered using scATAC-profiles and annotated using canonical marker genes, and the epithelial cell population subset was isolated (**Figure 6A, B**, **Supplemental Figure 2A**). Analysis of opened chromatin regions in bleomycin injured AT2 found increased gene activity scores of several abnormal genes including those associated with airway epithelial cells, i.e. *Scgb1a1*, *Muc5b*, *Sox2* and AT1/ transitional cell markers *Hopx* and *Krt19,* while the AT2 associated transcription factor *Etv5* was reduced (**Supplemental Figure 2B, C**) in YT^active^ bleomycin injured AT2 cells compared to WT bleomycin treated lungs. Marker gene analysis using promoter activity demonstrated that genes associated with airway (*Sox2, Runx2*), transitional cells (*Hopx, Krt19*), and proliferation (*Mki67*, *Ccna2*) were associated with AT2 cells from bleomycin injured YT^active^ mice (**Figure 6C**). Analysis of transcription factor binding site enrichment within opened promoter regions were associated with, among others, the AT2 cell regulator *Cebpa* (**Figure 6D**).

**Figure 6.**
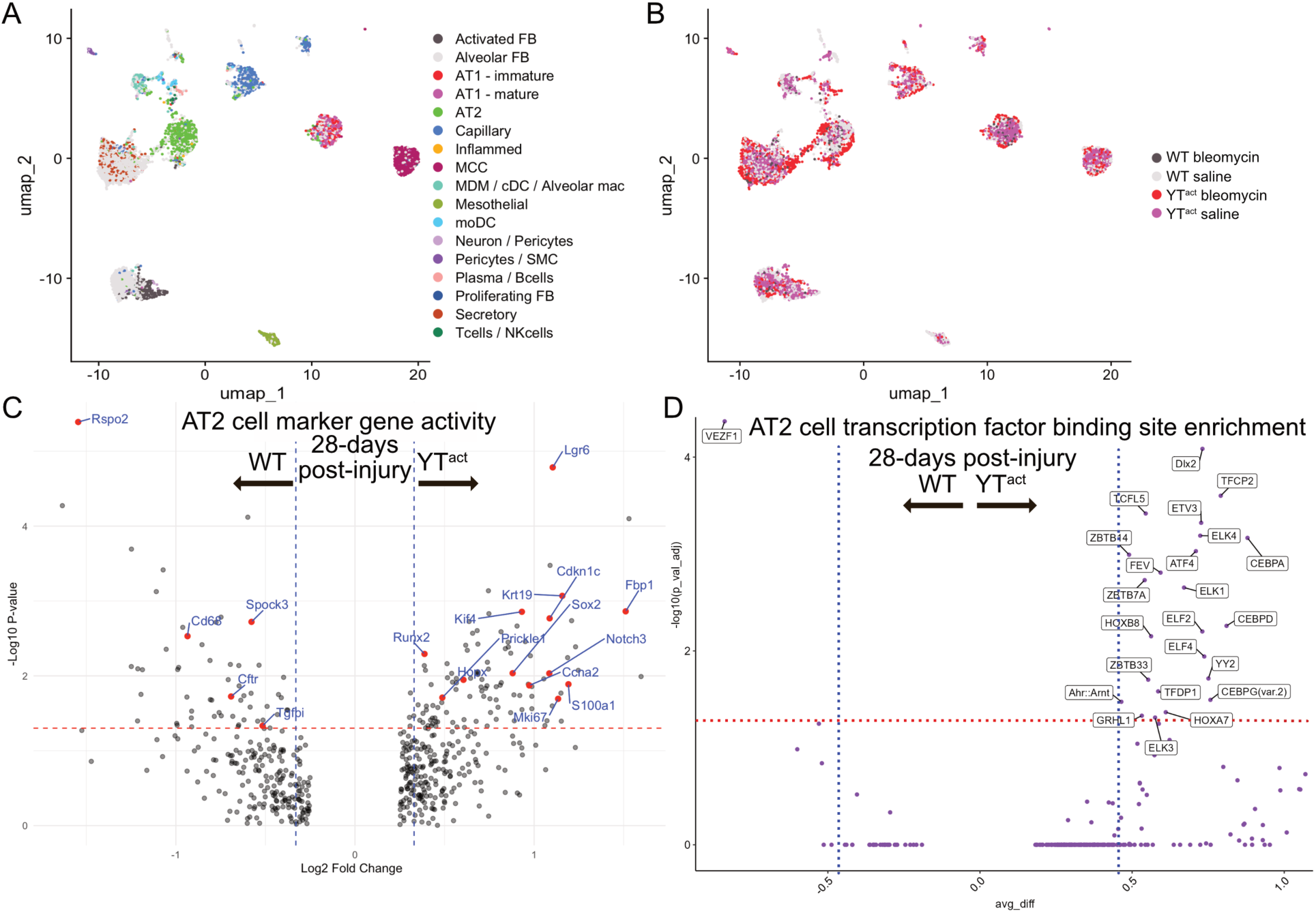
Single-Cell ATACseq demonstrates altered AT2 cell chromatin accessibility in YT^active^ bleomycin injured mouse lungs associated with aberrant transitional cells. **A**) UMAPs demonstrating clustering from 17 cell types in mouse lungs and (**B**) identifying cells recovered from respective genotype and treatment groups from snMultiome (RNA^+^ATAC-sequencing) of lung tissue from 28 days post-bleomycin. **C**) Volcano plot showing upregulated gene activity from AT2 cells of bleomycin injured wild-type (left) and YT^active^ (right) mice. **D**) Single cell ATAC sequencing shows enriched binding sites for open chromatin regions in bleomycin YT^active^ AT2 cells. The “avg_diff” is the fold-change computing the average difference in chromVAR z-score after performing differential testing between WT (left) and YT^active^ (right) mice.

To determine whether Verteporfin treatment corrected the abnormal AT2 differentiation identified by scATACseq, immunofluorescence and RNAscope analysis of lineage traced AT2 cells was used. RNA-ISH confirmed there were increased lineage-labeled *Krt19^+^* cells (12.4^+^/- 3.4% *Krt19^+^*/Lin^+^) in YT^active^ bleomycin injured mice compared to WT bleomycin injured lungs (0.9^+^/-0.4% *Krt19^+^*/Lin^+^). Verteporfin treated YT^active^ lungs had significantly reduced *Krt19^+^* cells compared to vehicle treated YT^active^ bleomycin injured lungs (**Figure 7A, B**) (7.7^+^/-1.5% *Krt19^+^/Lin^+^),* although this remained higher than bleomycin-injured Verteporfin treated WT mice (1.6^+^/-0.3%). With evidence of altered Cebpa activity from our snATAC-seq results, we then assessed a CEBPA gene module score from human AT2 cells grown in SFFFM (from Figure 2). Genes associated with CEBPA activity were reduced in human epithelial cells cultured with sustained Yap/Taz activation by XMU-MP-1 (**Supplemental Figure 2D**). Finally, immunofluorescence analysis of Cebpa in lineage traced AT2 cells from WT (91.0^+^/-3.6%) and YT^active^ (87.1^+^/-5.3%) mouse lungs treated with saline or WT bleomycin treated (89.9^+^/-6.0%) demonstrate that Cebpa is readily detected in lineage traced Sp-C^+^ AT2 cells, however this was significantly reduced (58.6^+^/-9.9%) in AT2 cells in YT^active^ mouse lungs injured with bleomycin. Verteporfin treatment significantly increased Cebpa^+^ lineage-traced cells in the YT^active^ bleomycin injured lungs (85.1^+^/-4.8%) (**Figure 7C, D**). Collectively these findings suggest sustained Yap/Taz activity negatively regulates Cebpa or that the sustained Yap/Taz activation leads to a loss of AT2 cell identity thereby reducing Cebpa expression. Treatment of Verteporfin improved alveolar repair including decreasing *Krt19^+^* transitional cells as well as restoring Cebpa expression in lineage traced AT2 cells. These findings indicate that disruption of aberrant persistent Yap/Taz activity can promote adaptive repair and potentially delay or stop the progression of pulmonary fibrosis.

**Figure 7.**
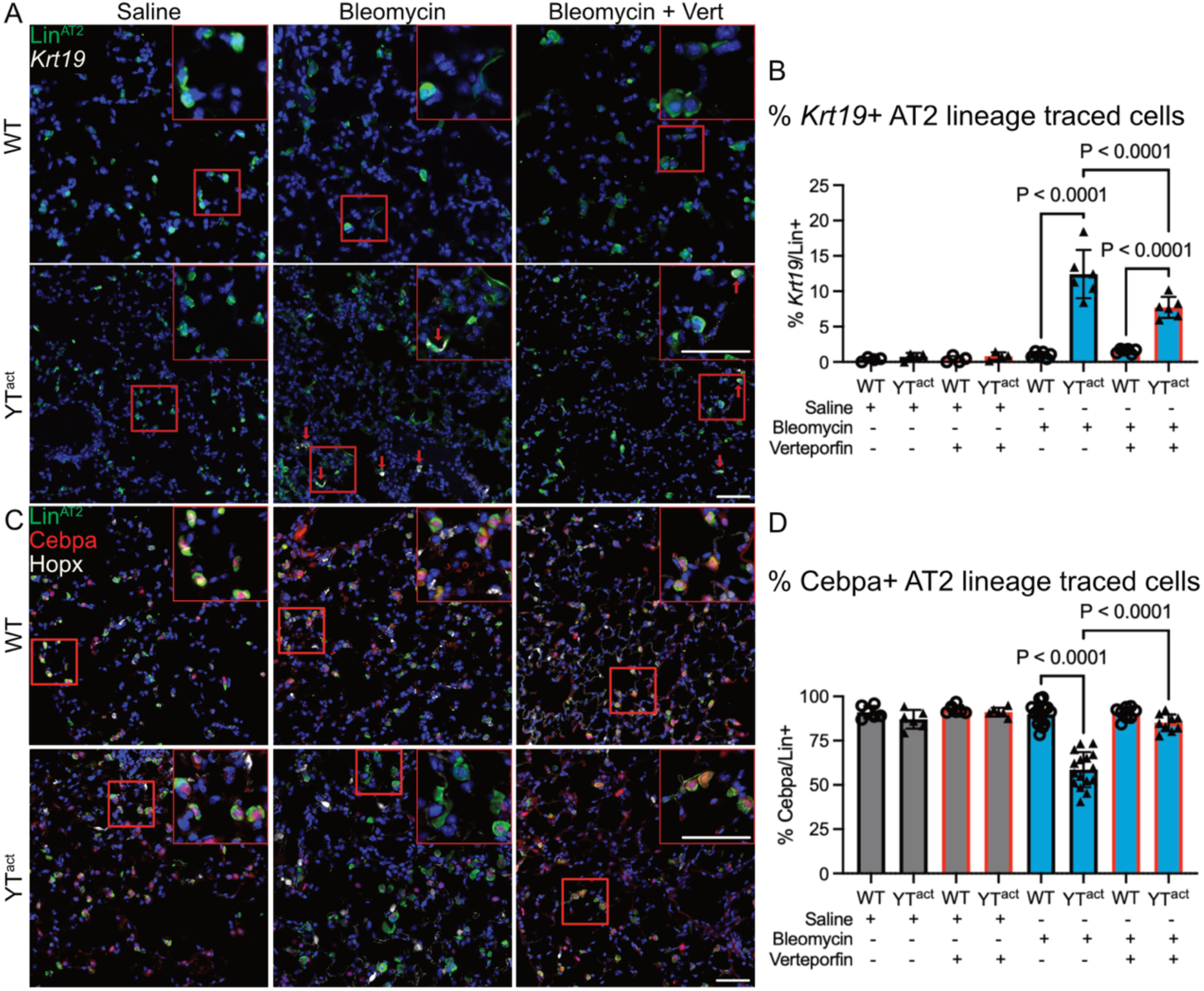
Sustained Yap/Taz activation leads to aberrant alveolar epithelial cell differentiation and accumulation of transitional cell populations which is partially corrected by Verteporfin treatment. **A**) Immunofluorescence and RNAscope analysis of *Krt19* (white) and lineage traced AT2 cells (green). **B)** Quantification of *Krt19^+^*/Lineage traced cells in each treatment group. WT and YT^active^ saline (n=4), WT and YT^active^ saline/Verteporfin (n=4), WT and YT^active^ bleomycin (n=6), and WT and YT^active^ bleomycin/Verteporfin-treated (n=6) mice. **C**) Immunofluorescence analysis of AT2 lineage-labeled cells (green), Cebpa (red), and Hopx^+^ AT1 cells (white) in wild-type and YT^active^ mice given saline, bleomycin, or bleomycin and Verteporfin. **D**) Quantification of Cebpa^+^ nuclei in AT2 lineage-labeled cells. Ordinary one-way ANOVA with Sidak’s multiple comparisons test with a single pooled variance was used to determine significance. Scale bars are 50μm.

## Discussion

Dysregulated repair of the alveolar epithelium is a hallmark of IPF. In this study, we demonstrate that both nuclear YAP and TAZ are increased in SP-C^+^ cells in the IPF lung. This is consistent with our previous work that showed increased YAP activity and loss of MST1/2 in the IPF epithelium^20,40,41^ and others have shown increased YAP in activated fibroblast populations^39,42,44,45,56,57^. Sustained YAP activity has also been identified in other forms of PF including Hermansky-Pudlak Syndrome^58^. Using a human alveolar organoid model, we found that pharmacologic activation of YT is sufficient to drive AT2 cells into a spectrum of aberrant “transitional” cell morphologies/phenotypes (but provocatively, not transcriptionally identifiable AT1 cells). Further, we found that YT activation in AT2 cells worsens experimental lung fibrosis and prevents repair, which could largely be ameliorated through treatment with a YT-inhibitor (Verteporfin) even after initiation of injury. Taken together, these results indicate that persistent YT activation in AT2 cells is maladaptive, preventing functional lung repair, and provide additional support for the concept that YT inhibition may be a promising therapeutic approach for IPF in the distal lung.

There have been several prior studies focusing on the role of Yap/Taz in lung development and injury repair using conditional Yap/Taz deletion strategies, while to date there has been limited investigation of the long-term impacts of persistent Yap/Taz activation in the lung epithelium^22,35–37,59,60^. In the developing lung, combined deletion of Yap/Taz using the Sftpc-CreERT2 and Sftpc-rtTA/tetOcre drivers led to reduced numbers of AT1 and AT2 cells^22,23^. Our current study, in concordance with our previous work showed Yap was rarely detected in AT1 cells and only expressed in AT2 cells during repair, while Taz was expressed in AT2 cells during repair and was readily detectable in most AT1 cells both during repair and at homeostasis^35,37^. Several studies have demonstrated that Yap/Taz are essential for AT2 to AT1 differentiation^22,23,61,62^ and are required for normal repair. Deletion of Yap/Taz prior to bleomycin, LPS, or bacterial injury results in worsened repair and fibrosis^35–37^. Others have shown that deletion of AT2 cell Taz alone is sufficient to prevent AT2 to AT1 differentiation and increased fibrotic response^38^. In addition, deletion of both Yap and Taz in Hopx^+^ cells resulted in loss of AT1 programming and caused these cells to adopt AT2-like features^59^. These studies lead to a paradigm in which YAP/TAZ is initially required for AT2 cell proliferation, while downregulation of YAP/TAZ supports AT2 cell maturation. Without initial YAP/TAZ activity, there is an absence of AT1 transitional cells and that at least TAZ is necessary to drive AT1 cell maturation and TAZ and/or YAP are required for AT1 cell maintenance. This concept shows a need for a finely regulated cell-type specific balance of YAP/TAZ activity during the repair process. Collectively, these findings demonstrate that activation of Yap and/or Taz is initially essential to guide adaptive alveolar epithelial repair, yet sustained aberrant AT2 activation promotes fibrotic remodeling.

Here, we find evidence that tight regulation of Yap/Taz activity is essential for proper alveolar repair. We previously showed that Yap activity in AT2 cells was increased early after bleomycin injury with peak nuclear localization at day 7, while Taz reached peak AT2 expression at 14-days post-injury. Both Yap and Taz returned to near zero activity by 21 days post-injury, indicating Yap/Taz are dynamically regulated during repair^35^. Our current study demonstrates that sustained activation of Yap/Taz in mouse AT2 cells did not alter the initial injury response or impact fibrosis at 14-days post injury but rather led to exacerbated fibrosis and the presence of persistent transitional cells by 28-days post-bleomycin that failed to resolve fibrosis by 8 weeks after injury. Inhibition of this sustained Yap/Taz activity with a single dose of Verteporfin at 14-days was sufficient to enhance fibrotic resolution in Yap/Taz active mice, although had no effect on WT mice at this time-point, further implicating that this was an epithelial Yap/Taz driven process. We acknowledge that this observation adds complexity to the growing body of literature in this area, including recent work that implied Hippo components inhibit alveolar repair (i.e., Yap/Taz activation may drive adaptive repair)^63^. There are several potential explanations for these seemingly opposing findings including differences in the degree and duration of Yap/Taz inhibition, as well as potentially distinct roles and timing of inhibition/activation of Yap and Taz in the repair process^37^. However, our results are consistent with the concept that there is sustained Yap/Taz activation in IPF and that sustained activation in human epithelial cells leads to abnormal epithelial cell differentiation.

Further, our in-vitro studies demonstrated that Yap/Taz activation was sufficient to lead to aberrant differentiation of human alveolar epithelial cells into KRT high and basal-like populations consistent with those seen in IPF^15,16,20^. Likewise, Yap/Taz activation resulted in increased *Krt19*^+^ transitional cells in mice following bleomycin injury. While the role of *Krt19* is not yet fully elucidated, these intermediate filaments appear as a convergent marker of epithelial cell states associated with injury and maladaptive/fibrotic remodeling^64^. This resulted in increased fibrotic remodeling of the mouse lung consistent with a concept that the presence of abnormal persistent transitional cells leads to fibrotic remodeling and the progression of pulmonary fibrosis^65,66^. These transitional cells, whether they initially occur during adaptive repair or are disease emergent, exist in multiple sub-types identified by various Krt markers (Krt8^+^, KRT17^+^/, KRT5^-^, Krt19^+^) or by various names (DAPTs, PATs)^46,67–71^. The presence of aberrant transitional cells has been identified in multiple forms of pulmonary fibrosis, including fibrosis associated with Hermansky-Pudlak Syndrome^71^, spontaneous fibrosis models associated with surfactant protein C mutations^66^, as well as bleomycin induced models of fibrosis^72,73^ and in IPF^15–17,74^. Our work has demonstrated that inhibition of the sustained Yap/Taz activation via Verteporfin resolves the presence of these Krt19^+^ transitional cells that were identified in the mouse lung, which coincided with decreased fibrotic remodeling. Recent work by others demonstrated that removal of other transitional cells, for example Krt8^+^ cells, also resulted in reduced fibrosis^65^. Collectively these studies demonstrate that the sustained presence of transitional cell populations, whether from normally occurring transitional cell populations or failed signaling pathways leading to aberrant differentiation of disease emergent populations, are associated with fibrotic progression and appear responsive to dynamic regulation of Yap/Taz activity.

This work also demonstrates that Yap/Taz sustained activation resulted in opened chromatin associated with Cebpa binding sites. We and others have demonstrated that Yap/Taz activation leads to opened/altered chromatin accessibility in the lung^22^ and other organs including the heart^75^. Generally, opened chromatin/increased presence of transcription factor binding sites are associated with increased activity. However, our immunofluorescent staining of Cebpa in lineage-traced AT2 cells, as well as gene module score in XMU-MP1 treated (YAP/TAZ activated via MST1/2 inhibition) human epithelial cells demonstrated decreased nuclear Cebpa localization and decreased gene module score respectively. These findings imply that Yap/Taz activity negatively regulates Cebpa. Cebpa was recently shown to be a regulator of AT2 cell fate and maintenance, as activation of Cebpa maintains AT2 cell transcriptional programs, while inhibition of Cebpa is necessary to allow AT2 cell differentiation into AT1 during both development and repair^76,77^. These studies implicate that there may be an inverse relationship between Yap/Taz and Cebpa in which there is a balance between Cebpa associated AT2 regulation and AT1 cell Yap/Taz associated signaling that regulates alveolar cell fate. These studies are consistent with our findings that Yap/Taz activation reduced Cebpa and recent work showing reduced alveolar Cebpa in the IPF lung^78^, and we have shown Yap/Taz to be activated in IPF. Inhibition of Yap/Taz via Verteporfin restored Cebpa nuclear localization in lineage traced AT2 cells, further supporting the concept that Cebpa and Yap/Taz signaling networks counter regulate each other.

One limitation of this study is that it does not completely discern whether the effect of Verteporfin is mediated predominantly by blocking the sustained activation of Yap/Taz in AT2 cells or both the sustained AT2 cell activation as well as the previously shown activation of Yap/Taz in activated fibroblast^39,42,44,57^. While others have reported different findings with Verteporfin treatment, our data indicate at minimum that the effects in the epithelium are important contributors to fibrotic remodeling. It is possible that some of the effects observed when treating human AECs in a reductionist organoid model with XMU-MP1 may differ from those seen in-vivo in which signaling from other cell types may intersect with this pathway. Finally, we acknowledge there may be differential timing or cell-type specific roles of Yap and Taz during alveolar differentiation and repair which will require further study.

Cumulatively, our findings demonstrate that the sustained activation of Yap/Taz via deletion of Hippo components Stk3/4 in mouse or by inhibition of the human homologues MST1/2 leads to the aberrant AT2 differentiation and persistence of transitional cells. This sustained Yap/Taz activation results in decreased fibrotic resolution resulting in persistent fibrosis 8-weeks after bleomycin induced lung injury. A single dose of Verteporfin was sufficient to promote resolution of transitional cell fates indicating restored alveolar repair and enhanced resolution of fibrosis in mouse injury models. These findings are consistent with the concept that in fibrotic models where persistent alveolar Yap/Taz activity are present, attenuating this pathway promotes resolution of fibrotic remodeling and potentially provides a therapeutic target to promote alveolar repair in pulmonary fibrosis.

## Methods

### Sex as a biological variable

To assess sex as a biological variable we used equal numbers of male and female mice for all studies. Secondary analysis found no significant difference between male and female mice and therefore for all subsequent analysis both male and female mice were combined for analyses.

### Human Subjects and Samples

Formalin-fixed, paraffin-embedded (FFPE) sections and fresh lung tissue for cell isolation and organoid culture was obtained from deidentified IPF and declined donor lungs removed at the time of lung transplant surgery from the lung tissue repository at Vanderbilt University Medical Center (IRB#060165, #192004).

### Human scRNA-sequencing reanalysis

Previously published scRNA-sequencing from IPF/ILD and control lungs (GSE227136) was reanalyzed for differential expression of genes in AT2 cells. AT2 cells (including proliferating AT2) from IPF and control samples were subset from the published integrated dataset and subject-level pseudobulk differential expression testing was performing using deSEQ2^79^.

### Human Organoids

Samples of human donor lungs were dissociated using dispase II (Roche 04942078001), collagenase I (Sigma-Aldrich C0130), and DNase (Millipore Sigma 260913-10MU) in phenol-free DMEM (Gibco 31053028). Cell suspensions were generated in C tubes (Miltenyi Biotec 130-093-237) using a gentleMACS dissociator then passed through 100 µm and 70µm filters (MTC Bio C4100 and C4070) to achieve single-cell suspensions. Epithelial cells were isolated by incubating cells with CD326^+^ microbeads (Miltenyi Biotec 130-061-101) and passing through LS columns (Miltenyi Biotec 130-042-401) positioned on a magnetic stand. CD326^+^ cells were plated in Matrigel droplets (Corning 356231) with serum-free feeder-free media (SFFFM) and expanded. Organoids were passaged in bulk, as whole organoids were liberated from Matrigel with 2mg/mL dispase for 1 hour with gentle disruption and passaged at 1:4 in Matrigel: SFFFM. At initial plating and after each passage Rock inhibitor was added to the media for days 1-3. Organoids were then treated with 3µm XMU-MP-1 (Tocris 6482) starting at day 4 for Yap/Taz activation. After 1 week, media was switched to either SFFFM or differentiation media (ADM) for another week then organoids were collected for end-point analysis.

### Human organoid cell morphology

Organoids generated from declined donor lungs were grown to confluency in domed Matrigel culture in SFFF media (StemCell Alveolar Organoid Media # 100-0847) then liberated as whole organoids with dispase (2mg/ml for 1hr with gentle disruption after 30min). Organoids were then counted and plated in 10ul of 1:1 Matrigel: SFFFM at ∼350 organoids per well into 96 well µ-Plate angiogenesis imaging plates (Ibidi 89646). Organoids were then cultured for 7 days in SFFFM with indicated treatments at day 4 (DMSO or 3µM XMU-MP-1 (MedChemExpress Cat. No.: HY-100526) then transitioned to Alveolar Differentiation Media (StemCell # 100-0861) or maintained in SFFFM for 7 days. Samples were then washed with PBS and fixed with 0.4% Glutaraldehyde for 20min at RT. Following 2x washes with PBS, 0.2% NaBH_4_ in dH_2_O was applied and incubated at 4C for 1hr for background quenching of autofluorescence of the Matrigel. Wells were then washed 3x in PBS. A staining solution in PBS-X (0.1% Triton X-100) containing Hoechst (15ug/ml), Syto14 (7.5 uM), Wheat Germ agglutinin conjugated to Alexafluor 555, (3.5ug/ml), and Phalloidin-568 (ThermoFisher A12380) (132nM) was prepared fresh and applied to the wells overnight at 4C with light excluded. Wells were then washed 3x for 40 minutes with PBS-X and imaged on an ImageXpress.AI (Molecular devices) as maximum intensity projections of 5µm Z-stacks across a depth of 40µm. All images were collected on the same day under the same settings for the entire plate. Images were analyzed using InCarta (Molecular Devices) with a SINAP custom trained algorithm for organoid segmentation based on TxRed (Phalloidin). Following organoid level masking (parent), nuclei were then segmented and analyzed. Standard multivariate analytic parameters within InCarta were used for quantification of organoids across all channels. Nuclei were only analyzed for morphology and RNA characteristics (SYTO14). Data was then analyzed in R using a custom pipeline starting at field level data (13 fields per well) consisting of metadata assignment, robust normalization followed by median absolute deviation (MAD) calculation and outlier removal in PCA space following Z-score calculation. Any missing data was then imputed with feature median, and columns with zero variance (failed metrics) were removed. Finally, PCA analysis was performed for display and phenotypic distance analysis in PCA space using Hotelling’s T-Test and multiple comparisons adjusted.

### Human organoid single-cell isolation and processing for scRNA-seq

To perform single-cell RNAseq, human organoids were treated in dispase (2mg/ml) for 1hr with gentle disruption after 30 minutes to break up the Matrigel and organoids. Organoids were washed twice in PBS and spun down at 500g. Organoids were then treated with 1mL trypsin (0.05%) for 7 minutes, then washed in PBS containing 5% BSA, washed 2 more times in PBS, then passed through 100μm, then 70μm cell filters to generate single cell suspensions. Samples were pooled across donors per treatment group. We then used 10X genomics single-cell RNA sequencing to target 10,000 cells per group, with 50,000 reads per cell. Donors were genetically demultiplexed using Demuxafy^80^ and the Vireo pipeline^81^. Clustering was performed and identified with marker genes using Seurat/Signac piepline^82,83^.

### Animal husbandry and YAP/TAZ gene regulation

These studies were approved by the IACUC at Vanderbilt University Medical Center. SftpcCreert2(blh)Stk3fl/flStk4*^fl/fl^* mice (JAX:028054^84^ crossed with JAX:017635^85^) were crossed with *Stk3^fl/fl^Stk4^fl/fl^*mice to generate experimental cohorts (YT^active^) with Cre^-^ littermates used as wild-type controls. SftpcCre*^ert^*^2r^*^osatdTomato^ Stk3^fl/fl^Stk4^fl/fl^*mice crossed with *Stk3^fl/fl^Stk4^fl/fl^* mice served as experimental cohorts for lineage-traced studies, with *S*ftpcCre*^ert^*^2r^*^osatdTomato^* mice (JAX:028054^84^ crossed with JAX:007909^86^) were used as lineage traced WT controls. Mice were administered tamoxifen (Millipore Sigma T5648) dissolved in corn oil via intraperitoneal (i.p.) injection at 100mg/kg either 3-weeks prior to or 2-weeks after bleomycin injury to induce Yap/Taz activation and lineage tracing. For Yap/Taz inhibition experiments, Verteporfin (Cell Signaling 64260) dissolved in DMSO was administered via i.p. injection at 60mg/kg 2-weeks after injury.

### Lung Injury and collection

Bleomycin (0.08 IU) was suspended in sterile saline and administered intratracheally (i.t.) at 100µL volume with equal volumes of saline administered as controls. Mice were sacrificed at indicated timepoints. The right lung was inflation-fixed with 10% buffered formalin at 25cm H_2_O pressure, excised, and submerged in 10% buffered formalin. The left lung was flash frozen in liquid nitrogen and processed for Sircol collagen analysis or for nuclei isolation for analysis.

### IHC Analysis

Inflation fixed mouse lungs (as described above) and organoids were fixed overnight at 4°C then processed and embedded in paraffin. Human lung samples were collected from de-identified transplant lungs from IPF patients and donor lungs collected were rejected for transplant. All histological staining was done on 5μm sections, and immunofluorescence and/or RNAscope RNA-ISH was conducted with antibodies and RNA probes listed in supplemental table 2. Citrate Buffer (pH 6, Sigma Aldrich, C9999) were used for antigen retrieval then slides were washed with deionized water. Slides were covered in block buffer (5% BSA) for 1 hour at room temperature. Block was removed and primary antibodies were added and allowed to incubate overnight at 4°C. Slides were washed 3X in PBST and secondary antibody was added and allowed to incubate for 2 hours at room temperature. For human lung samples, Vector TrueVIEW (Vector Laboratories Cat# SP-8400) was added as per the manufacturer’s instructions for 5 minutes to quench autofluorescence of red blood cells and collagen. Slides were washed again 3X in PBST and cover slips were applied with mounting media. Slides were imaged capturing 10 non-overlapping frames/lung with a 20X objective on a Keyence BZ-X710 inverted fluorescence microscope.

### Fibrosis Scoring

Semi-quantitative analysis of fibrosis was done utilizing modified Ashcroft scoring (Hübner et al., 2008) of Masson’s trichrome stained slides. For each lung, 2 independent blinded scorers assessed at least 10 20X objective magnification frames/lung and scores were averaged. If there was a scoring discrepancy of greater than 1.5, scores were discussed, and consensus was reached. Collagen content was quantified using Biocolor’s sircol soluble collagen assay kit (Biocolor S5000).

### Single-cell Multiome Sequencing and Analysis

Single nuclei multiome sequencing (ATAC^+^RNA) was performed using 10x genomics chromium sequencing targeting 10,000 nuclei per sample 50,000 reads per nuclei. Single Nuclei were isolated from flash frozen left lung tissue pooling 1 male and 1 female mouse per group and sequencing was aligned to GRcm39 and total of 21,144 nuclei were sequenced post processing across samples. Analysis was performed by the Creative Data Solutions genomic analysis group at Vanderbilt University Medical Center. Gene expression mapped to the genome with relatively low confidence/read depth. SoupX and scDoublet were used to remove ambient DNA and detect doublets. Clustering was performed and identified with marker genes using Seurat/Signac piepline^82,83^ using scRNA sequencing from other in-house mouse lung samples to identify cell-type specific clusters which resulted in 17 cell types identified. ATACseq was then mapped to these clusters with open chromatin associated with marker genes used to map to clusters identified from the RNA expression analysis using Harmony. Plots were made using ggplot2 and scCustomize. For ATACseq analysis MACS2^87^ was used for peak calling. Gene activity scores were generated using peaks within 2kb upstream of the transcriptional start sites. Transcription factor enrichment within promoters were identified using Chromvar enriched motif with Signac^88^.

### Statistics

Statistics were performed using Graphpad Prism 10. Normality was tested using Shapiro-Wilk and Kolmogorov-Smirnov tests. For 2 variable comparisons, unpaired parametric t-tests were used. For comparisons with more than 2 variables, ordinary one-way ANOVA with Sidak’s multiple comparisons test with a single pooled variance was used.

### Study Approval

Mouse studies were approved by Vanderbilt University Medical Center’s IACUC, under approval M1500027 and M2400059. Deidentified human lung samples were acquired, and human studies were approved under protocol 060165 and 192004 by Vanderbilt University Medical Center’s human subjects IRB.

## Data Availability

Genomic data is available for download at GSEAXXXX (Data are being uploaded to the Gene Expression Omnibus and analysis script is available at Github https://github.com/KropskiLab/Gokey-2025_yaptaz_activation).

## Author Contributions

IPG, ASM, JAK, and JJG designed research studies. IPG, ASM, NMG, ACC, GTD, TPS, UKS, HED, APS, and JJG conducted experiments. IPG, ASM, NMG, GTD, TPS, DSN, and JJG acquired data. IPG, ASM, ACC, JPC, SS, SSG, TSB, JAK, and JJG analyzed data. IPG, ASM, JAK, and JJG wrote the manuscript. All authors edited to the manuscript

## Acknowledgements

The authors meet criteria for authorship as recommended by the International Committee of Medical Journal Editors (ICMJE) and were fully responsible for all aspects of the study and publication development. This was supported by NIH/NHLBI R01HL145272 (JAK), R01HL153246(JAK) Vanderbilt Faculty Research Scholars (JJG, ASM), The Francis Family Foundation (JJG, ASM), R01HL151016 (TSB), The Pulmonary Fibrosis Foundation Fellowship (ASM), The Vanderbilt Institute for Clinical Translational Research (VICTR) VR55627 and VR55246 (JJG) under CTSA 5UL1TR002243. Imaging obtained for morphologic analysis were captured on the ImageXpress Micro Confocal High Content Screening System which is housed and managed within the Vanderbilt High-Throughput Screening Core Facility, an institutionally supported core, and was funded by NIH Shared Instrumentation Grant 1S10OD028719.

---

**Supplemental Table 1:** Patient data including age and sex of donor lungs rejected for transplant and IPF lungs used for analysis of YAP/TAZ expression and organoids associated with Figures 1-3.

| Sample | Lung state | Age | Smoking Status | COD | Notes |
| --- | --- | --- | --- | --- | --- |
| VUHD84 | Healthy Donor | 34/Female | 1p/d/7 years | CPA |  |
| VUHD85 | Healthy Donor | 53/Male | none | CVA/Stoke | Hypertension, Family History of CAD |
| VUHD87 | Healthy Donor | 53/Male | 1p/d/7 years | Cardiac Arrest | Etoh Abuse, Hypertension, Family history of CAD |
| VUHD92 | Healthy Donor | 55/Male | none | CVA/Stroke | Hypertension, Family history of CAD |
| VUHD94 | Healthy Donor | 62/Male | unknown | Stroke | No medical history |
| VUHD140 | Healthy Donor | 61/Male | unknown | CVA | No medical history |
| VUILD72 | IPF | 62/Female | None |  | HTN,DM2, Dyslipidemia, GERD, Osteoarthritis, Rheumatoid Arthritis, Restless Leg Syndrome, Moderate Asthma |
| VUILD74 | IPF | 62/Male | 10 PY quit 40 years ago |  |  |
| VUILD76 | IPF | 53/Male | none |  | Obesity, Hyperlipidema |
| VUILD77 | IPF | 62/Male | 30 PY quit 18 years ago |  | DM2, HTN, Hypothyroidism |
| VUILD79 | IPF | 59/Female | Quit 25 years ago |  | GERD, HLN |
| VUILD80 | IPF | 69/Female | N/A |  | No medical history |

**Supplemental Table 2:** Antibodies and RNAscope probes used, along with catalog number, supplier, and concentrations used.

| Antibody | Target Species | Company | Catalog # | Concentration used |
| --- | --- | --- | --- | --- |
| Ager | Rat | R&D Systems | 175410 | 1:200 |
| Alexa Fluor Conjugated HOPX647 | NA | Santa Cruz Biotechnology | sc-398703 | 1:100 |
| Alexa Fluor Conjugated CCSP488 | NA | Santa Cruz Biotechnology | sc-365992 | 1:100 |
| GFP | Chicken | Rockland Immunochemicals | 600-901-215 | 1:300 |
| TROMA-I (Krt8) | Mouse | DSHB | TROMA-I | 1:300 |
| SPC | Rabbit | Abcam | AB90716 | 1:100 |
| TdTomato | Goat | SICGEN | AB8181-200 | 1:500 |
| RNA Scope | Mouse | ACD Bio | Mm-Krt19-C2 | 1:100 |

**Supplemental Figure S1:**
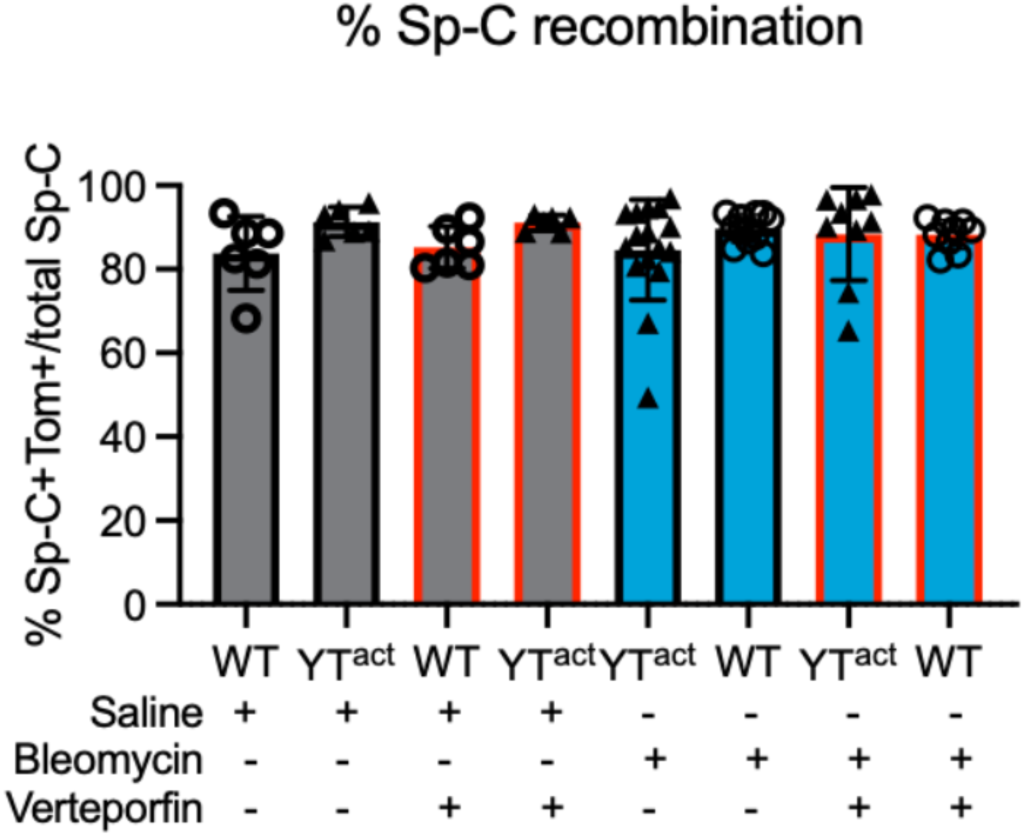
Quantification of Sp-C^+^/Lineage traced cells to determine recombination efficiency of the mouse model. Supplemental figure is associated with main Figure 4.

**Supplemental Figure S2:**
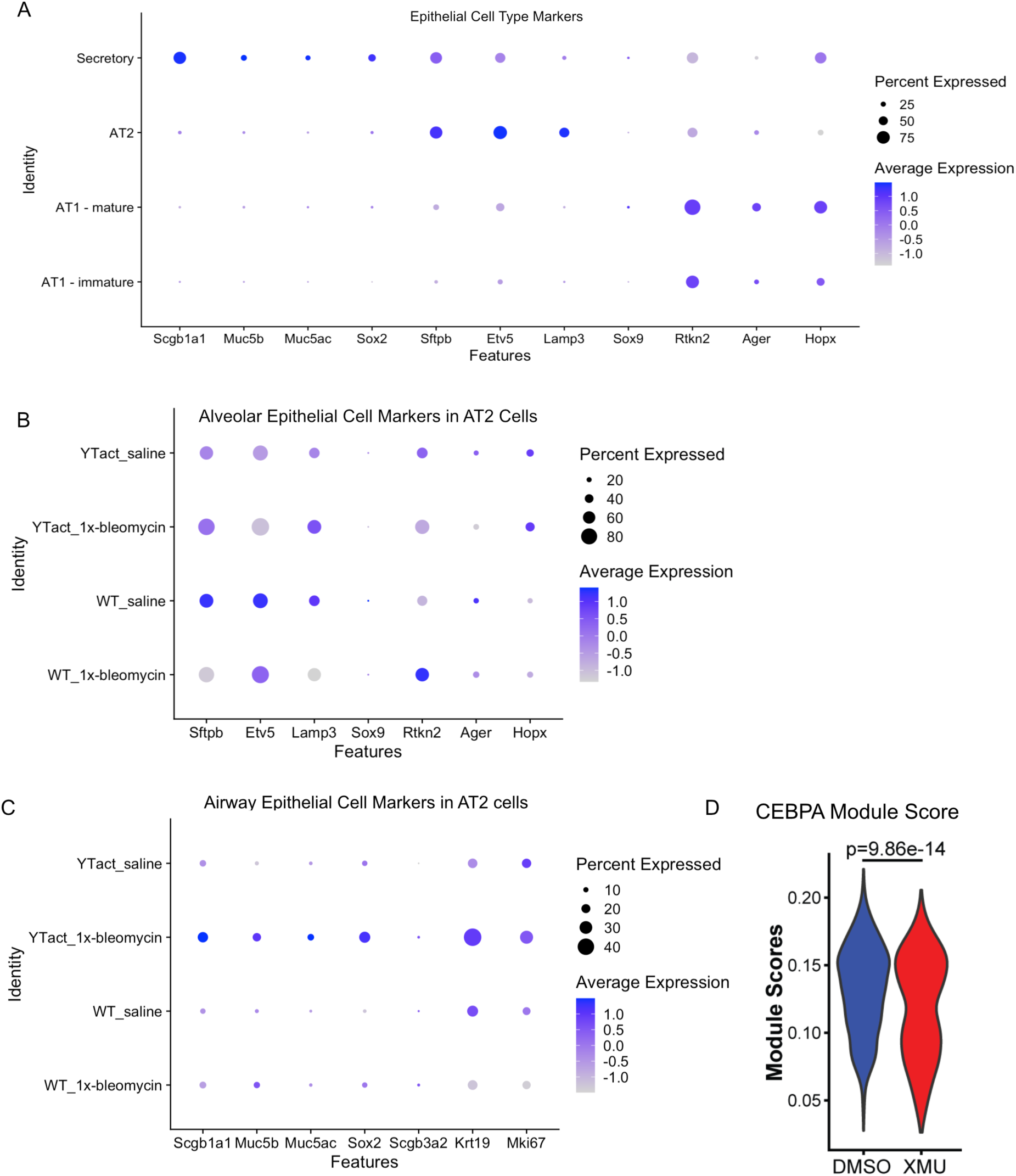
**A)** Dotplot of cell type markers of the 4 epithelial cell types identified by our single-nucleus multiome (RNA^+^ ATACseq) analysis associated with Figure 5. **B**) Dotplot showing AT2 cell associated gene activity scores and **C**) Dotplot showing airway, transitional, and proliferation marker gene activity scores determined from single-cell ATAC sequencing associated with Figure 5. **D**) Gene module score showing decreased signaling associated with CEBPA activity from scRNAseq of human organoids from Figure 2.

